# DarHTG is a tripartite phage defence system for purine DNA base ADP-ribosylation

**DOI:** 10.64898/2026.11.02.763450

**Authors:** Raphael Ledermann, Anna-Lena Gotzler, Barbara Steigenberger, Yijun Wang, Johanna Zimmermann, Selina Geiger, Jieying Fu, Johannes Hoffmann, Osamu Suyari, YuGeng Zhang, Yang Lu, Andreja Mikoč, Christoph Tang, Ivan Ahel, Marion Schuller

## Abstract

Bacteria evolved a variety of phage defence strategies to combat the constant onslaught of viral threats. DarTG toxin-antitoxin systems provide phage defence by ADP-ribosylation of bases in single-stranded DNA which interferes with viral replication and aborts phage infection. Here, we describe DarHTG, a system widespread among bacteria, that combines DarTG with a non- processive superfamily I helicase functioning as co-toxin for DarT and thereby enabling the ADP- ribosylation of double-stranded DNA. We identified two such DarHTG subclasses which catalyse the ADP-ribosylation of adenosine and guanosine bases, respectively, and reverse these modifications with structurally unique ARH superfamily antitoxins. Thus, DarHTG antitoxins expand the known macromolecular target spectrum of the ARH enzyme class to nucleic acids. These DarHTG systems confer resistance to a broad range of bacteriophages and can be triggered by the highly charged tails of viral single-stranded DNA-binding proteins, which occlude the active site of the ARH antitoxin.

## Introduction

Nucleic acid modifications represent a widespread strategy involved in bacterial defence against phage. DNA modifications can serve as means of self-versus-non-self-discrimination, *e.g.* in restriction- modification systems, or can be targeted by bacterial defence systems which recognise chemically modified bases such as 7-deazaguanine or 5-hydroxymethylcytosine that are often incorporated into phage DNA to shield against bacterial nucleases^1–4^. Furthermore, ADP-ribosylation has been identified as a strategy for bacterial immunity by physically blocking phage replication, transcription and translation machineries^5–8^. Conserved in all domains of life and viruses, ADP-ribosylation is a reversible modification of small- and macromolecules^9,10^. The diverse enzyme superfamily of ADP- ribosyltransferases (ARTs) catalyses the ADP-ribosylation reaction transferring ADP-ribose (ADPr) from *β*-NAD^+^ onto target substrates^11^. ADP-ribosylation is regulated by a dynamic interplay between ARTs and ADP-ribosylglycohydrolases removing ADPr modifications^12^, which include three evolutionary unrelated protein families, *i.e*. macrodomains, NADARs, and ARHs^13–15^. The first phage defence system discovered to utilise DNA ADP-ribosylation is the toxin-antitoxin (TA) system DarTG^5^. In this system, a <u>D</u>NA <u>A</u>DP-ribosyltransferase (DarT) toxin catalyses the ADPr modification, while the <u>D</u>NA <u>A</u>DP-ribosyl<u>g</u>lycohydrolase (DarG) antitoxin reverses it. Thus far, two DarTG systems have been described, with DarTG1 and DarTG2 shown to mediate reversible ADP-ribosylation of guanosine (DarTG1)^13^ or thymidine (DarTG2)^16^ bases in single-stranded DNA (ssDNA). Whilst the DarT proteins are structurally similar between both systems, a different DarG ADP-ribosylglycohydrolase superfamily protein is associated with the two systems, *i.e.* NADARs (‘DarG_N_’) in DarTG1^13^ and macrodomain- containing proteins (‘DarG_M_’) in DarTG2^5,16^. DNA ADP-ribosylation is highly toxic to the bacterial cell, resulting in cell death^5,17,18^. In non-infection conditions, suppression of DarT toxicity by its cognate antitoxin is essential for the survival of bacteria. Phage infection, however, results in DNA ADP- ribosylation, inhibiting phage DNA replication followed by abortive infection^6^.

Here, we describe DarHTG, a tripartite TA system that combines DarTG with a non-processive superfamily I (SF1) helicase, ‘DarH’^19^. We define DarHTG-associated helicases as co-toxins for DarT that facilitate the ADP-ribosylation of double-stranded DNA (dsDNA). We identified two DarHTG subclasses that catalyse the ADP-ribosylation of adenosine and guanosine bases, respectively, thus establishing DarHTG as a system for purine DNA base ADP-ribosylation. Furthermore, DarHTG encodes DarG antitoxins belonging to the ARH superfamily which counteract helicase-DarT-induced toxicity by reversing the DNA ADP-ribosylation, thereby expanding the known macromolecular targets of these enzymes from proteins to nucleic acids. DarHTG systems are widespread among bacteria and confer resistance to a broad range of bacteriophages. We show that DarHTG activity can be triggered by the highly charged tails of ssDNA-binding proteins of phages occluding the active site of the ARH antitoxin, which provides the first insight of a molecular trigger of a DarTG-based phage defence system.

## Results

### DarHTG is a tripartite toxin-antitoxin phage defence system dependent on an SF1 helicase

Whilst the established DarTG systems are canonical two-component TA systems^5,20^, DarHTG combines *darT* and *darG* with an SF1 helicase (‘DarH’) encoded upstream within the operon (Fig. 1a and b). Phylogenetic analyses identified two clades of DarT toxins, ‘DarT3’ and ‘DarT4’, associated with DarHTG systems and which cluster close to but separate from DarT1 toxins (Fig. 1c). DarHTG systems are widespread among prokaryotes, and found in *e.g. Escherichia*, *Bacillus*, *Burkholderia*, *Vibrio,* and *Mycobacteria* species, and associated human pathogens. We cloned different *darHTG* operons representing the two DarHTG subclasses. While expression of *darT1* or *darT2* by themselves is highly toxic^5,13^, DarHTG-associated DarT toxins showed substantial toxicity only when both, *darT* and *darH* were co-expressed (Fig. 1d), establishing DarH as a crucial mediator of toxicity. The ARH protein (‘DarG_A_’), encoded by *darG_A_* within the *darHTG* operon acted as the antitoxin of DarHTG systems (Fig. 1d). We purified DarT3 and DarH from *Escherichia coli*, and tested their ADP-ribosylation activity on purified genomic DNA (gDNA). In agreement with the observed toxicity mediated by DarHT, only the presence of both proteins led to ADP-ribosylated gDNA (Fig. 1e). Modification was abolished, when either *β*-NAD^+^ (the source of ADPr), or ATP or Mg^2+^ (known cofactors of SF1 helicases) were omitted from the reaction mixture (Fig. 1e). Furthermore, *E. coli* constitutively expressing DarHTG from *Trichlorobacter ammonificans* showed resistance against phage Maverick (‘Mav’)^21^, with strict requirement for *darH* (Fig. 1f). Moreover, such infection resulted in the accumulation of ADP-ribosylated DNA (Fig. 1g). As was reported for DarTG1 and DarTG2 systems, DarHTG likewise provides phage defence via abortive infection. Mav suspension spotted onto a lawn of *darHTG*-expressing or control *E. coli* caused plaques (indicating initial infection of both) which increased in diameter only in the absence of the DarHTG system, indicating that no functional phage particles were produced in presence of DarHTG, a hallmark of abortive infection (Fig. 1h).

**Figure 1.**
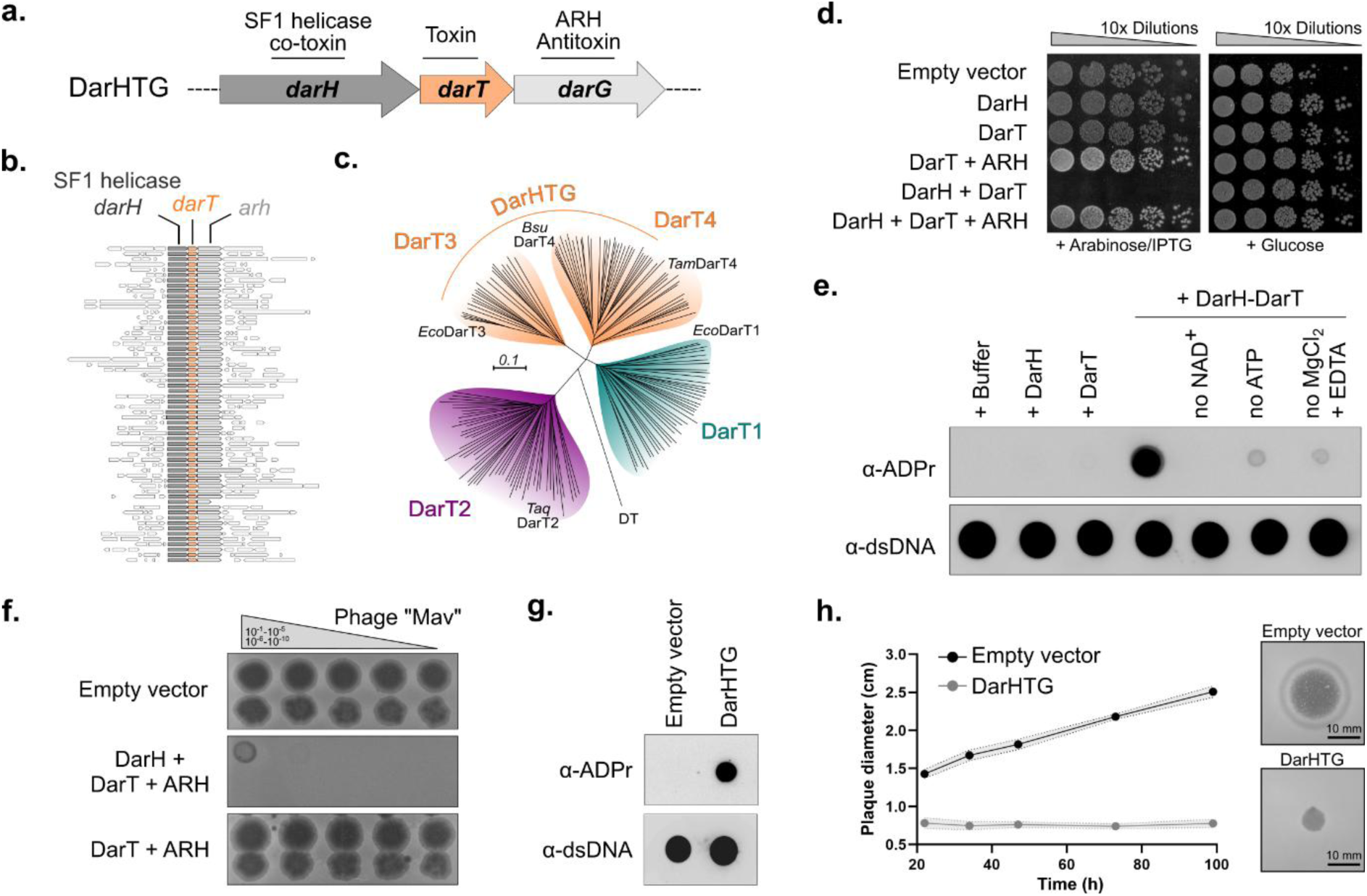
The phage defence system DarHTG. **a.** DarHTG is a tripartite toxin-antitoxin system, characterised by a DarT toxin which is flanked by a SF1-helicase (DarH) upstream and an ARH enzyme (DarG) downstream. **b.** Gene neighbourhood analysis showing conserved genetic association of DarT homologues with genes coding for DarH and ARH (n=60). **c.** DarT toxins encoded within a DarHTG operon context, termed DarT3 and DarT4, cluster phylogenetically different to DarT1 and DarT2. Rooted with diphtheria toxin (DT). Representative DarTs for each cluster are highlighted. **d.** DarH functions as co-toxin and ARH as antitoxin for DarT activity. Toxicity assay monitoring the growth of *E. coli* BL21(DE3) under repression (glucose) and induction (arabinose/IPTG) of the expression of *B subtilis darHTG* components is shown. **e**. Immunodot blot showing the *in vitro* ADP-ribosylation of genomic DNA (gDNA) by *E. coli* DarH and DarT which requires the co-factors *β*-NAD^+^ for DarT and ATP/MgCl_2_ for DarH. **f)** *T. ammonificans* DarHTG protects against phage infection with DarH being required for its function. **g)** *T. ammonificans* DarHTG induces ADP-ribosylation of DNA upon infection with phage Mav compared to vector control. Samples were analysed by immunodot blotting 40 min after infection. **h)** DarHTG phage defence relies on induction of abortive infection. Quantification of plaque diameter size (Data are mean ± s.d.; *N* = 4 per time point), showing steady increase over the course of infection in *E. coli* carrying the empty vector control compared to *T. ammonificans* DarHTG- expressing cells.

### DarH is an atypical SF1 helicase enabling DarHTG to target dsDNA

FoldSeek^22^ revealed the replicative SF1A helicases^23^, UvrD and Rep, as the closest homologues of DarH. Indeed, AlphaFold3^24^ models of DarH closely superimpose with the ATP-binding RecA-like domains 1A and 2A, and the DNA-coordinating domain 1B of UvrD (r.m.s.d. = 4.0 Å), however, DarH lacks domain 2B present in both UvrD and Rep (Fig. 2a). Comparison of the enzymatic activity of purified *E. coli* DarH with UvrD showed that both are functional ATPases, with their activity stimulated by DNA, as previously observed for UvrD^25^. However, UvrD was around 100-fold more active compared to DarH (Fig. 2b). Furthermore, whereas UvrD showed the expected high dsDNA unwinding activity^26^, DarH activity was very limited, indicating DarH does not function as a processive helicase (Fig. 2c). The presence of the cognate DarT slightly stimulated DarH unwinding activity by approximately 2-fold, although this remained far below the activity of the processive helicase UvrD (Fig. 2c). Helicases are dependent on both NTP hydrolysis and divalent cations for their activity^23^, and the same was true for DarH (Figs. 1e, 2d), albeit with a relaxed co-factor specificity. While SF1 helicase members show a preference for ATP^23^, DarH accepted ATP, GTP, CTP, UTP and even deoxy nucleotides (dNTP mix) *in vitro*. Cation testing showed that DarH accepts Mg^2+^, Ca^2+^ and Mn^2+^, but would tolerate neither Co^2+^ nor Zn^2+^ (Fig. 2d) as was found for other SF1 helicases^27^. We further tested the specificity of DarHTG-associated helicases for DarHT-induced toxicity. Indeed, *B. subtilis* DarT strictly depended on its cognate DarH. Neither *T. ammonificans* nor *E. coli* DarH complemented the absence of the cognate DarH, and neither could UvrD or Rep, even following deletion of their 2B domains (Fig. 2e). While DarT toxins are generally characterised to modify ssDNA, we hypothesised that the presence of and dependency on DarH allows DarHTG systems to also target dsDNA. We performed *in vitro* gel-shift ADP-ribosylation assays using a hairpin structure with a high predicted T_m_ of 98°C, thus resulting in a dsDNA stretch inaccessible for DarT toxins. Indeed, *E. coli* DarT3 alone could not modify this substrate, however, multiple ADPr modifications were obtained when DarH was present (Fig. 2f). Notably, *E. coli* DarT3 was able to ADP-ribosylate the top strand when provided isolated as an ssDNA substrate without the presence of DarH, showing that DarH can generate ssDNA substrates *in situ* from dsDNA sequences for DarT (Fig. 2f).

**Figure 2.**
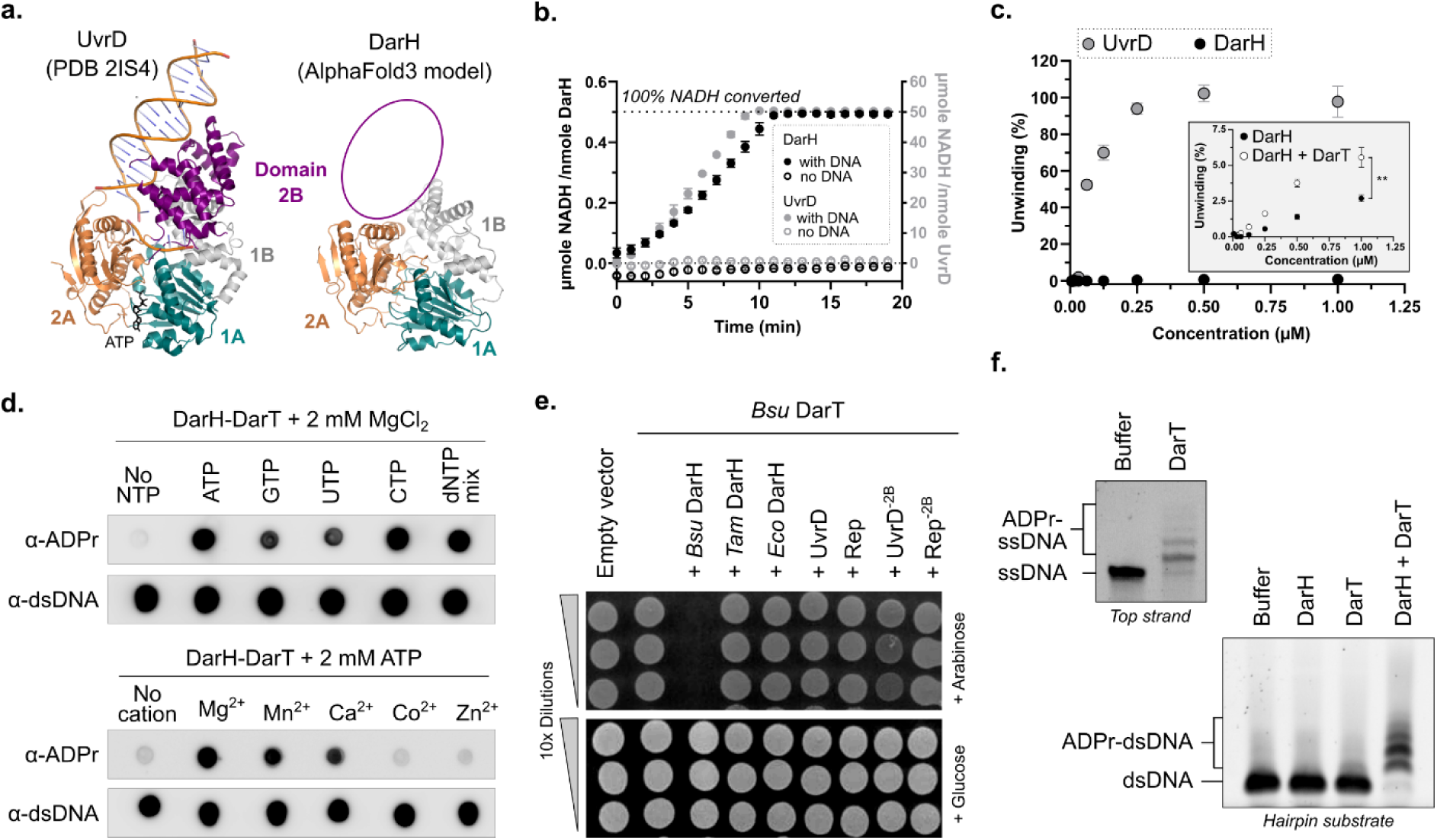
DarHTG-associated helicases are UvrD-like helicases facilitating dsDNA ADP- ribosylation. **a.** Comparison of the crystal structure of *E. coli* UvrD (PDB ID 2IS4) with an AlphaFold3 model of *E. coli* DarH reveals structural conservation apart from domain 2B (purple). The conserved RecA domains (1A and 2A) are shown in orange and cyan. The three conserved core domains of *E. coli* UvrD and *E. coli* DarH superimposed with an r.m.s.d. of 4.0 Å. Same angle of view is shown. **b.** *E. coli* DarH shows ATPase activity upon DNA binding but with 100-fold less activity compared to *E. coli* UvrD. ATPase activity was measured using an LDH/pyruvate-based NADH conversion assay. Data are mean ± s.d.; *n* = 3. **c.** DarH possesses no DNA unwinding processivity compared to UvrD. The presence of DarT does not notably increase DNA processivity as shown in the inset of the graph. The main graph shows % of unwinding after 10 min, the inset shows % of unwinding after 60 min. Data are mean ± s.d.; *n* = 3, ** p-value of 0.0024 from unpaired, two-tailed t-test. **d.** DarH can utilise a variety of nucleotides while showing cation preferences. Immunodot blots showing the *in vitro* ADP-ribosylation of gDNA by *E. coli* DarH and DarT in presence of nucleotides and cations. **e.** Toxicity is induced only by DarH cognate to the co-expressed DarT. Toxicity is not induced by non-cognate DarHs as well as UvrD/Rep and variants lacking the 2B domain. Growth of *E. coli* BL21(DE3) is monitored under repression (glucose) and induction (arabinose) of the expression of DarT and helicases. **f.** DarH facilitates ADP-ribosylation of dsDNA shown by gel-shift assays. The model dsDNA substrate folds into a hairpin structure that is only modified by DarT in presence of DarH. The top strand of the hairpin structure is modified by DarT when provided as ssDNA.

### Two classes of DarHTG-associated DarT toxins ADP-ribosylate adenosine and guanosine DNA bases

We tested *E. coli* DarT3 and *B. subtilis* DarT4 ADP-ribosylation activity on PolyT ssDNA oligomers containing a single A, G, or C base insertion to test their base specificity. *E. coli* DarT3 modified only PolyT-A with a single modification, whereas *B. subtilis* DarT4 showed exclusive activity on PolyT-G (Fig. 3a). This base specificity of DarT3 and DarT4 for adenosine and guanosine bases, respectively, underlies the phylogenetic clustering into two distinct DarT clades (Fig. 1c). Neither enzyme modified the analogous ssRNA oligomers (PolyU-A and PolyU-G) (Fig. 3b). Since DarT2 toxins show a sequence motif specificity, we permutated the bases directly adjacent to the modified A and G, respectively, with other bases in the PolyT backbone. For both DarT toxins we found some sequence context dependency for the targeted base, with the most marked effect exercised by the base directly upstream. The preference for pyrimidine bases adjacent to the modified adenine was particularly evident for *E. coli* DarT3. Furthermore, both enzymes could only modify one of two subsequent As or Gs, respectively, potentially due to steric reasons (Fig. 3c).

**Figure 3.**
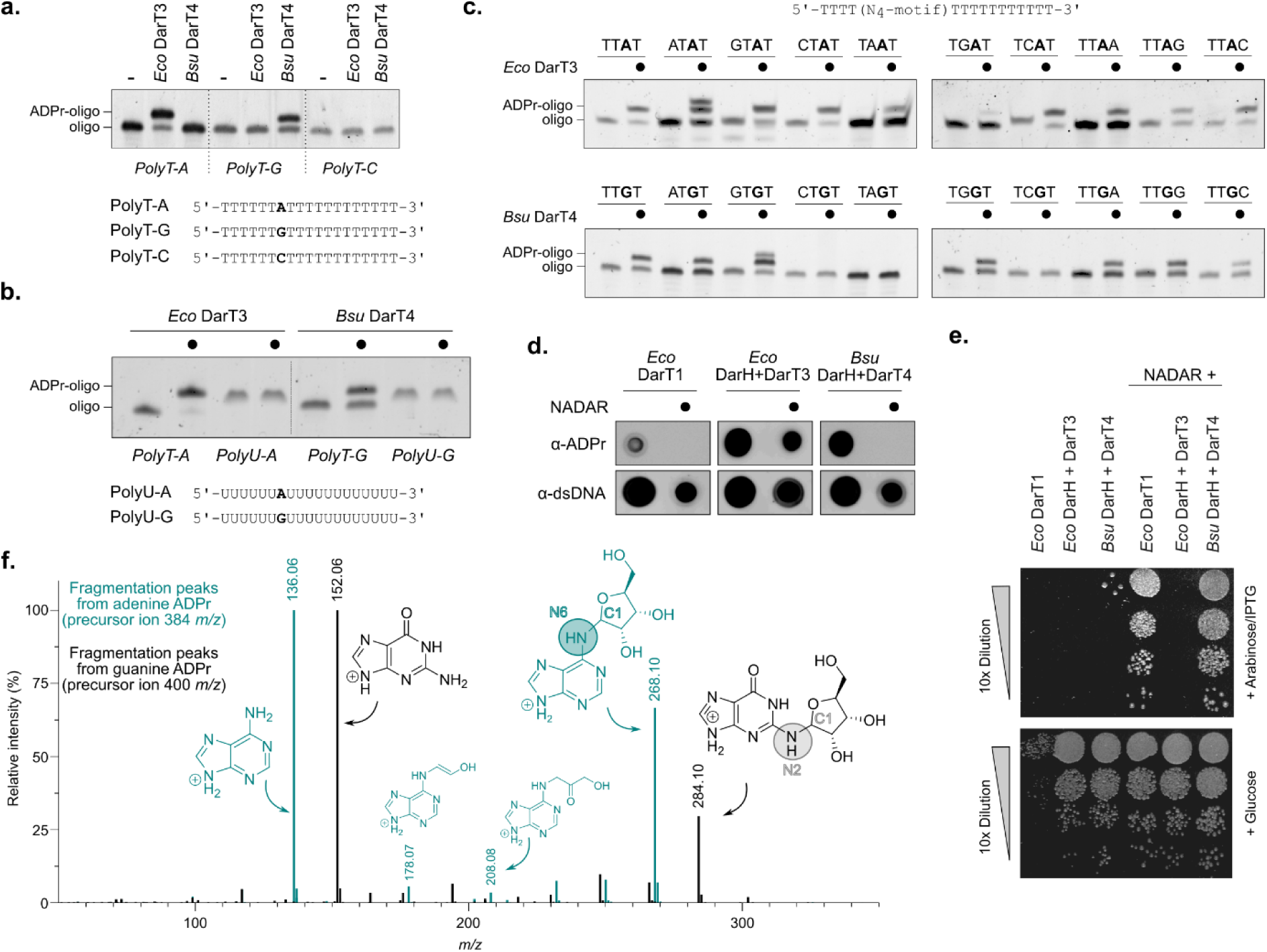
DarT3 and DarT4 toxins ADP-ribosylate adenosine and guanosine DNA bases. **a.** Gel- shift assay showing ADP-ribosylation activity of *E. coli* DarT3 and *B. subtilis* DarT4. **b.** DarT3 and DarT4 targets DNA and not RNA for ADP-ribosylation. **c.** DarHTG-associated toxins show DNA base preferences in proximity to the ADP-ribosylated adenosine (*E. coli* DarT3, top) or guanosine (*B. subtilis* DarT4, bottom) DNA base. **d**. DarG_N_ efficiently reverse *B. subtilis* DarH-DarT4-catalysed guanosine ADP-ribosylation in contrast to *E. coli* DarH-DarT3-catalysed adenosine ADP-ribosylation. **e.** DarG_N_ rescue from *B. subtilis* DarH-DarT4-induced toxicity but not from *E. coli* DarH-DarT3 induced toxicity. Toxicity assay is shown, monitoring the growth of *E. coli* BL21(DE3) under repression (glucose) and induction (arabinose/IPTG) of the expression of *darG_N_*, *darH*, *darT3* and *darT4* enzymes. **f.** Overlay of MS/MS fragmentation spectra of ADP-ribosylated-adenosine product catalysed by *E. coli* DarH-DarT3 (cyan) and ADP-ribosylated guanosine product catalysed by *B. subtilis* DarH-DarT4 (black). Predicted fragments are depicted with molecular structures next to the *m/z* peaks.

Adenosine bases present a new DNA base target for DarT toxins, whereas ADP-ribosylation of guanosine bases was previously shown for DarT1 toxins^13^. Notably, DarG_N_ of *E. coli* DarTG1 efficiently reversed ADP-ribosylation of gDNA, isolated from cells expressing *B. subtilis darHT4*, but not from cells expressing *E. coli darHT3* (Fig. 3d). Likewise, DarG_N_ rescued from *B. subtilis* DarHT4-, but not from *E. coli* DarHT3*-*induced toxicity (Fig. 3e). In complementary gel-shift assays, (Fig. 4h), DarG_N_ did also not reverse *E. coli* DarHT3-ADP-ribosylated dsDNA, similar to *B. subtilis* DarG_A_ or *T. aquaticus* DarG_M_. To confirm these findings, indicating that DarT4 catalyses the same reaction as DarT1 while DarT3 establishes an ADPr-adenosine base linkage so far undescribed for DarTG-based systems, we performed LC-MS/MS analyses on ADP-ribosylated PolyT-A and PolyT-G oligos (Fig. S1a) as previously described^7,28^. ESI-MS/MS fragmentation of the [M+H]^+^ precursor ions for ribosylated adenosine and ribosylated guanosine at 384 and 400 *m/z,* respectively, resulted in the fragment ions for adenine (136 *m/z*) and guanine (152 *m/z*) but also for ribosylated-adenine (268 *m/z*) and ribosylated-guanine (284 *m/z*) (Fig. 3f). Additionally, we identified the fragment ions 178 and 208 *m*/*z* (Fig. 3f, S1b) which have ribose fragments (including the anomeric C1 of ADPr) attached to the adenine and which result from ADP-ribosylation of the adenine N6^7^. Furthermore, the acquisition of a matching reference spectrum (Fig. S1c, d) obtained from the PolyT-G substrate modified by *Streptomyces coelicolor* ScARP, which is known to attach the ADPr to the N2 of the guanine^29,30^, confirmed that *B. subtilis* DarT4 establishes the same linkage between the anomeric C1 of ADPr and the amino N2 of the guanosine base (Fig. 3f).

**Figure 4.**
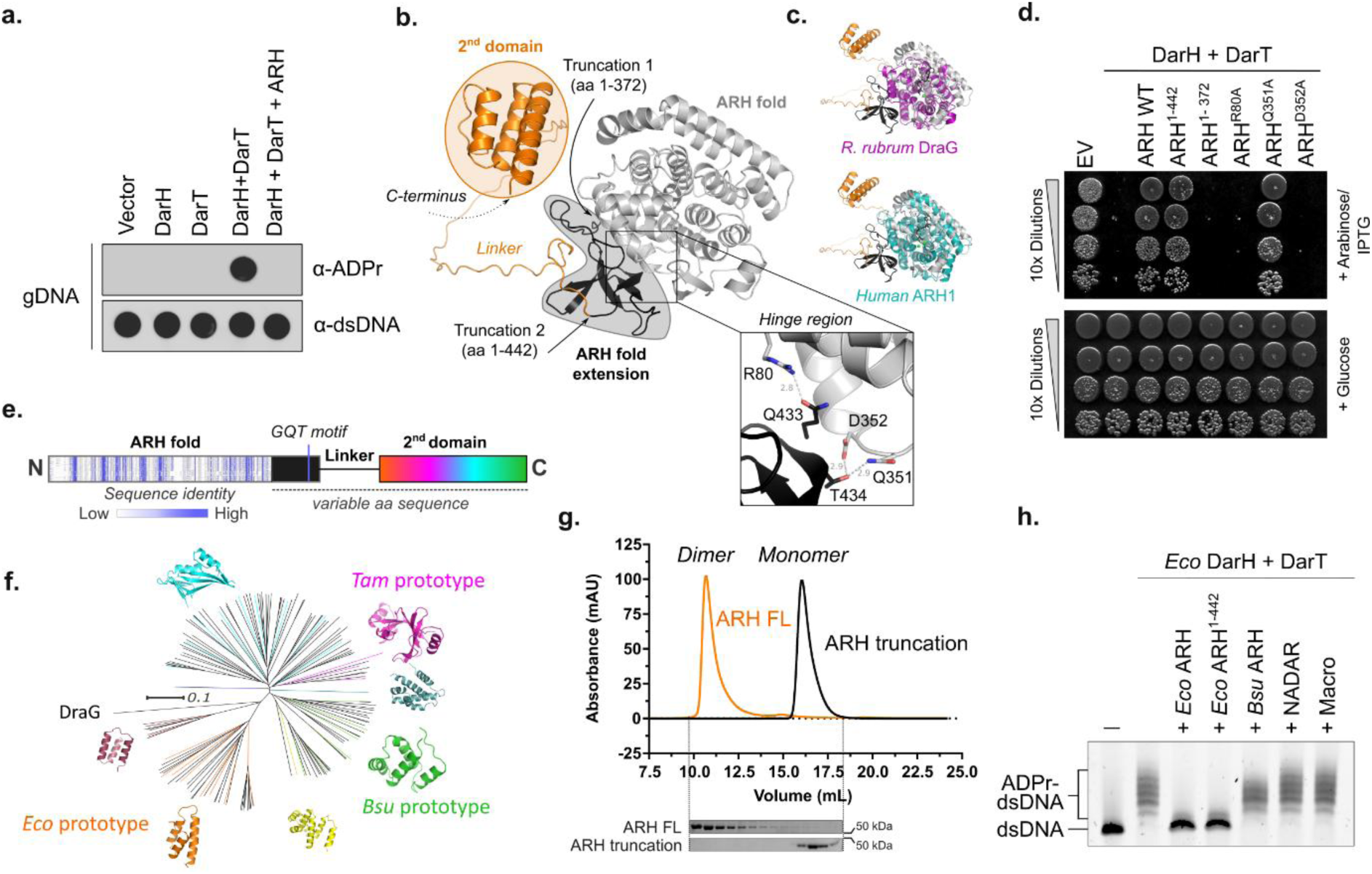
ARH enzymes with unique features function as DarHTG antitoxins. **a.** DarG_A_ enzymes reverse DarH-DarT-catalysed ADP-ribosylation. Immunodot blot of gDNA isolated from *E. coli* expressing *B. subtilis darHTG* components as indicated is shown. **b.** AlphaFold3 model of *E. coli* DarG_A_ antitoxin showing the ARH fold (grey), ARH fold extension (black) and C-terminal secondary domain attached to a linker (orange). **c.** The ARH fold extension (black) and secondary domain (orange) are unique compared to known protein-targeting ARH enzymes. Structural overlay of *E. coli* DarG_A_ with phylogenetically closest bacterial (*R. rubrum* DraG, r.m.s.d = 3.8 Å, PDB ID 2WOE) and human (ARH1, r.m.s.d = 4.5 Å, PDB ID 6G28) ARH homologues is shown. **d.** The secondary domain is not required for DarG_A_ antitoxin function. Toxicity assay is shown, monitoring the growth of *E. coli* BL21(DE3) under repression (glucose) and induction (arabinose/IPTG) of the expression of *E. coli darH*, *darT* and *arh* enzymes as indicated. **e.** Schematics of sequence alignment of DarG_A_ enzymes. Full sequence alignment is shown in the Supplementary Information Fig S2a. **f.** Phylogenetic groups of DarG_A_ homologues cluster based on their variable secondary domains. Coloured lines in tree indicate ARH variants modelled by AlphaFold3, revealing the secondary structure shown around the tree. **g.** Secondary domains of DarG_A_ function as dimerisation domains. Overlay of gel filtration profiles showing the elution of *E. coli* ARH full-length and ARH^1-442^ truncation, with SDS-PAGE of analysed fractions shown below. **h.** *E. coli* DarH-DarT dsDNA ADP-ribosylation of adenosine bases is reversed by the ARH fold of the cognate DarG_A_ and not by antitoxins of the guanosine-targeting DarHTG (*B. subtilis* ARH), DarTG1 (*E. coli* C7 NADAR) and DarTG2 (*T. aquaticus* macrodomain (macro)) systems.

### Structurally unique ARHs as antitoxins of DarHTG systems

Similar to NADARs of DarTG1 and macrodomains of DarTG2, ARH enzymes associated with DarHTG systems remove the ADP-ribosylation modifications on gDNA in bacteria, thus rationalising their antitoxin function for DarHTG systems (Fig. 4a). AlphaFold3^24^ models revealed unique structural features of DarG_A_ compared to characterised bacterial and eukaryotic ARHs including the closest homologues *Rhodospirillum rubrum* DraG and human ARH1 (Fig. 4b, c). DarG_A_ not only contains the characteristic ARH core fold, which closely superimposes with other ARHs, but also an ARH fold extension, which interacts with the ARH core via a hinge region, and an unstructured linker ending in a small C-terminal, secondary domain (Fig. 4b, c). In *E. coli* DarG_A_, the interaction of the ARH fold extension with the ARH fold is predicted to be stabilised by hydrogen bonds between R80 and Q433 and between Q351/D352 and T434, with Q433 and T434 being located within a conserved GQT motif of the ARH fold extension (Fig. 4b, S2a). The ARH fold extension and its stabilisation to the ARH fold via the hinge region is essential for DarG_A_ antitoxin function (Fig. 4d). Whilst the ARH fold contains highly conserved residues, including the expected ARH active site residues, both the extended core fold (apart from the GQT motif) and the C-terminal domain show very little primary sequence conservation (Fig. 4e, S2a). Notably, AlphaFold3^24^ models of the C-termini of various DarG_A_ showed large variances in the length of the unstructured linker, and also a plethora of different folds of the small C-terminal domain, resulting in different ARH clades (Fig. 4f). Even though members of each clade had a similar fold, the primary sequences within clades were poorly conserved (Fig. S2b). AlphaFold3^24^ predicted that each of these C-terminal domains promotes the formation of ARH dimers (Fig. S2c). Analytical size exclusion chromatography confirmed that the full length but not the truncated *E. coli* ARH indeed forms a dimer (Fig. 4g). However, toxicity assays showed that dimerisation was dispensable for antitoxin function *in vivo* (Fig. 4d), hydrolytic activity *in vitro* (Fig. 4h), or phage defence (Fig. S2d).

### DarHTG systems confer resistance to a broad range of bacteriophages

Screening of the DarHTG systems selected for this study against the BASEL I^31^, BASEL II^32^, and the Durham^21^ phage collections, revealed that both the *T. ammonificans* and *E. coli* DarHTG system protected *E. coli* strains from a broad range of different phages. The *E. coli* system conferred broader phage protection, with its phage specificity profile largely overlapping that of the narrower range of the *T. ammonificans* system (Fig. 5a and b). The *B. subtilis* DarHTG system was not protective against any of the 136 tested phages. Derived from a Gram-positive bacterium, it is unclear if the physiology of the host itself affected the system or the *E. coli* phages cannot be recognised by the system. Notably, we found that DarHTG systems from *T. ammonificans* (also a Proteobacterium) or even *E. coli*, were markedly affected in their functioning based on the expressing *E. coli* strain (Fig. S3).

**Figure 5.**
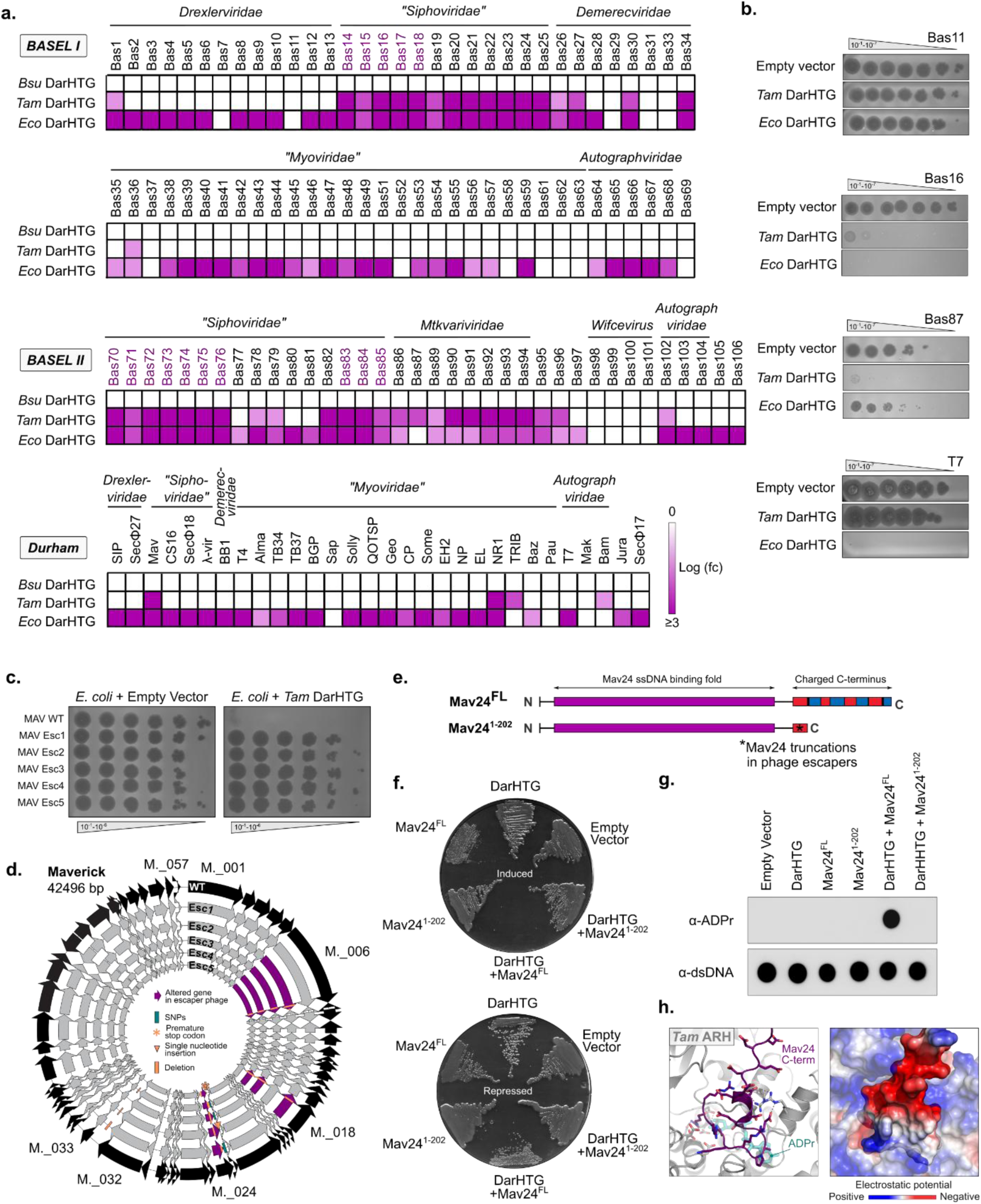
Phage defence by DarHTG systems. **a.** Phage specificity profiling of DarHTG from *B. subtilis*, *T. ammonificans*, and *E. coli*. Purple shading reflects 10-fold changes in pfu. Reduction in plaque size has been counted as an additional fold change if applicable. Phages of the BASEL collection highlighted in purple contain the gene DUF2815, *i.e.* referred to as Mav24 in the Mav phage. **b.** Representative examples of phage plaque assays are shown on the right, complete phage profiling data is shown in the Supplementary Information Figures S7-1 to S7-4. **c.** Phage plaque assay of Mav wild-type phage and Mav-derived phages (Esc1-5) escaping *T. ammonificans* DarHTG-mediated defence. **d.** Genome representation of Mav wild-type (WT) and escaper phages (Esc1-5) with mutations highlighted as indicated by the legend within the circle. **e.** Comparison of Mav24 (DUF2815) full-length with Mav24 truncations in escaper phages Esc1 and Esc3-5. **f.** Induction of co-expression of *T. ammonificans* DarHTG with Mav24 full-length results in toxicity in *E. coli* compared to co-expression with Mav24 truncation. **g.** Immunodot blot showing the presence of ADP-ribosylated DNA upon co-expression of *T. ammonificans* DarHTG with Mav24 full-length compared to co-expression with Mav24 truncation. **h.** AlphaFold3 complex model of *T. ammonificans* DarG_A_ with C-terminus of Mav24, showing occupation of the ADPr binding site. (Left) Cartoon stick model generated by overlay with ADPr ligand from co-crystal structure of ARH1 (PDB ID 6G28), (right) model with surface electrostatics mapped onto *T. ammonificans* DarG_A_ and Mav24 C-terminus.

### An ssDNA binding protein acts as a phage trigger for DarHTG

We noticed papillations on initial plaques of Mav on *T. ammonificans darHTG*-expressing cells (Fig. S4). We purified five individual plaques, and confirmed these as escaper phages unaffected by the *T. ammonificans* DarHTG system (Fig. 5c). Full-genome sequencing of the escaper mutants revealed multiple SNPs, but all five carried mutations leading to a similar C-terminal truncation of Maverick_024 (“Mav24”), which is annotated as DUF2815 (Fig. 5d, S5a). Structure predictions of Mav24 revealed an N-terminal ssDNA-binding domain and a large unstructured C-terminus (Fig. S5b), which contains four repeats of negatively-positively-charged stretches, thereby ending on a positively charged stretch (Fig. S5a,b). All escaper mutants had the core DUF2815 fold intact but carried truncations just after a negatively-charged stretch (Fig. 5e, S5a). Whilst high induction of full length and the truncated Mav24 versions were toxic (Fig. S5c), we noticed that weak induction only led to cell death in the presence of constitutively-expressed *T. ammonificans darHTG* in case of full length but not truncated Mav24 (Fig. 5f). Moreover, we detected the presence of ADP-ribosylated DNA exclusively in cells expressing both full-length Mav24 and *T. ammonificans darHTG* (Fig. 5g). Complex modelling between the C-terminus of Mav24 and *T. ammonificans* ARH predicted an interaction between the positive stretch of the Mav24 C-terminus with the active site of the ARH fold (Fig. 5h, S5d) which coordinates and compensates the charge of Mg^2+^ ions with negatively charged aspartate/glutamate side-chains^33^. Furthermore, *T. ammonificans* ARH shows positively charged patches around its active site, potentially involved in DNA recognition, which are predicted to interact with the negative stretch of the Mav24 C-terminus (Fig. 5h, S5d). These features rationalise the structural complex models which predict that the complementarily charged C-terminus of Mav24 targets and occludes these sites (Fig. 5h), thereby preventing ARH substrate binding and catalysis. A BLAST search of the Mav24 sequence against phages from the BASEL collections showed that 15 phages harbour Mav24 homologues; all of these being susceptible to the *T. ammonificans* DarHTG system (Fig. 5a, highlighted in purple). However, numerous Mav24-deficient phages were still susceptible to *T. ammonificans* DarHTG and the Mav escaper mutants isolated against *T. ammonificans* DarHTG were still susceptible to *E. coli* DarHTG (Fig. S5e). This suggests that these ssDNA-binding proteins represent only one of potentially several triggers for DarHTG activation.

## Discussion

Tripartite TA systems present a recently defined concept for anti-phage defence^19,34^ compared to canonical dual TA systems^35^. Incorporation of a third component, that serves as a chaperone, regulator, or an additional antitoxin or toxin, increases their functional diversity and regulatory potential^34^. DarHTG is a DarTG-based TA system which incorporates the UvrD-like SFI helicase DarH. DarH acts as the co-toxin of DarT, with DarT being dependent on DarH for induction of toxicity, DNA ADP- ribosylation, and phage defence. Compared to classically defined dual-toxin systems among the tripartite TA systems^34^ including KKP, which provides the two active toxin kinases^36^, DarH provides the substrate for DarT which then generates the toxic product. Griesman *et al.*’s studies suggest that DarH forms a complex with DarT which increases its affinity for DNA and thereby its ADP-ribosylation activity. Furthermore, our findings show that DarH allows the targeting of dsDNA for ADP-ribosylation by providing the DNA substrate as a single-strand required for the DarT ADP-ribosylation reaction. The DarH-DarT cooperativity therefore makes DarHTG the first DarTG-based phage defence system mediating dsDNA ADP-ribosylation.

DarH is structurally similar to UvrD-like helicases, which were also shown to be components of the PARIS and Gabija phage defence systems^37^. Unlike UvrD, however, DarH lacks the 2B domain, which mediates both the interaction of UvrD with the dsDNA duplex and UvrD dimerisation through a 2B– 2B interface^38,39^, with the latter being required for long distance unwinding^38,40^. Monomeric UvrD can still unwind a limited amount of DNA^40^ and with the co-localisation guided by the interaction between DarH and DarT proposed by Griesman *et al.*, long-distance unwinding of DNA by DarH may not be needed for its function in contrast to the house-keeping helicases UrvD and Rep. Local unwinding may be sufficient to provide DarT the substrate for ADP-ribosylation directly at the site targeted by DarH. Domain 2B is not required for general helicase activity^39^ as demonstrated by DarH retaining ATP hydrolysis activity upon DNA binding.

Two DarHTG-associated DarT clades were identified that differ in the purine base targeted for ADP- ribosylation. While DarT3 targets adenosine, DarT4 ADP-riboslyates guanosine DNA bases, both with a slight motif preference of TA and TG, respectively. Similar observations have been made by Griesman *et al*.. Guanosine base ADP-ribosylation at the N2 position has been characterised as the activity of DarT1 toxins and other eukaryotic and bacterial toxins including pierisins and ScARP^13,41,42^. However, DarT3 toxins are the first enzymes discovered that catalyse robustly and efficiently adenosine DNA base ADP-ribosylation *in vivo*. So far, only CmdT toxins of the phage defence system CmdTAC (a tripartite TA system with a chaperone as third component) was known to target adenosine bases efficiently, albeit within mRNA^7^. Notably, both enzymes, *i.e.* CmdT and DarT3, establish the same molecular linkage between the adenine N6 and the anomeric C1 of the NAM-proximal ribose of the ADPr. Thus, DarT toxins of both DarHTG subclasses attach the ADPr to the amino group of the purine DNA base, thereby catalysing an *N*-glycosidic linkage.

The antitoxins of both DarHTG subclasses are members of the ARH superfamily. DarG_A_ therefore complete the ‘trinity’ of known ADP-ribosylglycohydrolase superfamilies acting as antitoxins for DarTG-based systems. Moreover, DarHT3 and its associated ARH enzyme present the first system for specific and reversable ADP-ribosylation of adenosine DNA bases. The closest homologues of DarG_A_ are bacterial DraG and human ARH1, that are characterised to hydrolyse arginine ADP-ribosylation from proteins^15,43^. The homology becomes evident in the ability of DarG_A_ to hydrolyse an *N*-glycosidic linkage, analogous to the *N*-glycosidic linkage between ADPr and arginine hydrolysed by the protein- targeting ARHs. While catalytic mechanisms may therefore follow the same principles, DarG_A_ antitoxins expand the known macromolecular target spectrum of ARHs from proteins to nucleic acids, likely facilitated by their unique structural features additionally to the conserved ARH fold. Mapping surface electrostatics onto *B. subtilis* and *E. coli* DarG_A_ structure models revealed positive electrostatic surfaces generated by several basic residues in the ARH fold extension around the catalytic site (Fig. S2e), supporting DNA substrate recognition for ADPr-DNA hydrolysis. The function of the DarG_A_ dimerisation domains and the reason for their variability in sequence and structure remains to be clarified.

Directed evolution revealed that DarG_A_ targeting presents a way to activate DarHTG-mediated phage defence. All isolated escaper mutants, derived from the phage Mav and resistant to *T. ammonificans* DarHTG, showed truncations of Mav24, annotated as DUF2815 and predicted as an ssDNA-binding protein. Structural modelling suggested the complementarity of the highly charged C-terminus to the electrostatic charge on the surfaces around the catalytic site of *T. ammonificans* DarG_A_, allowing the C- terminus of Mav24 to occlude the active site of the antitoxin, which leads to uncontrolled DNA ADP- ribosylation by DarHT and ultimately, aborts the phage infection (Fig. 6). DUF2815 homologues can be found in genomes of phages and bacteria, but not in eukaryotes. Bacterial DUF2815 homologues generally have no charged, unstructured C-termini compared to viral homologues (Fig. S6). This indicates that DUF2815 is functional without the unstructured C-terminus consistent with the escaper phages still being infectious. Overall, Mav24 mutations illustrate an elegant strategy employed by escaper phages to evade defence by *T. ammonificans* DarHTG and give first insights into a possible phage trigger for a DarTG-based phage defence system.

**Figure 6.**
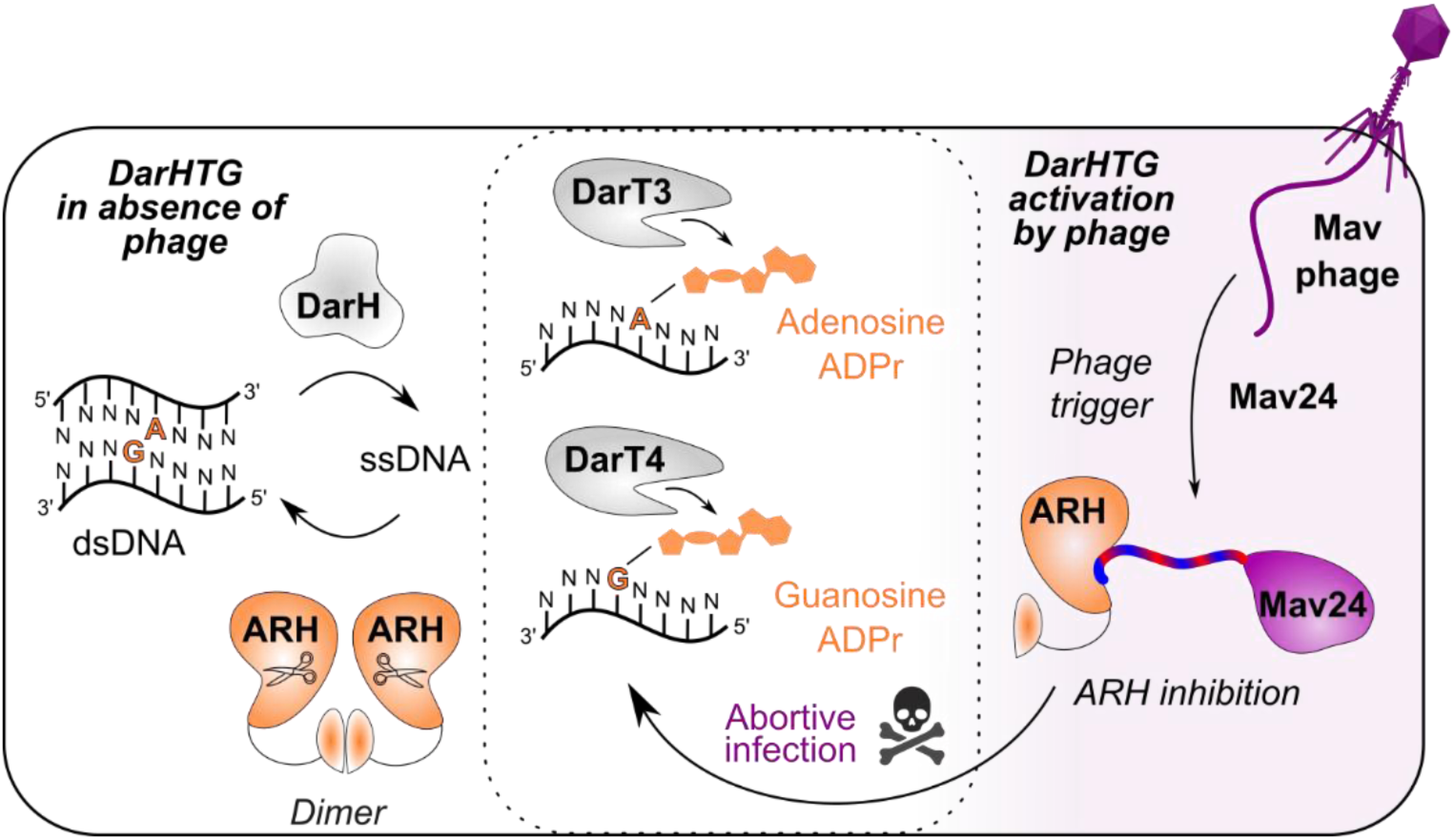
Model of DarHTG-mediated phage defence. *Please refer to discussion for details*.

Phage specificity profiling and the resistance of *E. coli* hosting a DarHTG from a different bacterial species towards Mav24 mutant phages indicates that Mav24 presents one of likely several DarHTG triggers. These observations are in line with the DarHTG family being widespread among prokaryotes and conferring resistance to a broad range of bacteriophages. This study provides functional insights into all three components of the DarHTG tripartite system and establishes a foundation for future studies to elucidate the mechanisms and triggers governing system activation, as well as its interplay with host factors and potentially other phage defence systems, which together shape the outcome of DarHTG- mediated phage defence.

## Materials and Methods

### Bacterial strains and culture conditions

*E. coli* strains were cultivated in lysogeny broth (LB) at 37°C with appropriate antibiotics included at the following final concentrations: ampicillin 100 µg/mL, chloramphenicol 17 µg/mL, kanamycin 50 µg/mL, streptomycin 50 µg/mL, and tetracycline 10 µg/mL. For routine maintenance, strains carrying genes encoding toxins under the control of the P_BAD_ promoter were grown in medium supplemented with 0.8% (^w^/_v_) glucose to repress promoter activity. Bacterial strains used in this study are listed in Supplementary Table S1.

### Strain and plasmid construction

To combat the high toxicity of DarHT of the *B. subtilis* and *T. ammonificans* DarHTG system for cloning, the NADAR sequence of *E. coli* C7^13^ was chromosomally integrated into *E. coli* DH5α (NEB) as described by St-Pierre *et al*.^44^. For this, the NADAR DNA sequence was cloned by Gibson assembly into the ‘One-Step Integration Plasmid’ (pOSIP) encoding a kanamycin resistance marker for integration at the phage P21 integration site. The integration module and antibiotic resistance marker, flanked by FRT sites, were removed using pE-FLP for FLP recombinase-mediated excision. Integration of *nadar* was verified by Sanger Sequencing, resulting in strain ‘DH5α-NADAR’. Cloning strategies of plasmids generated in this study are described in detail in the method section of the Supplementary Information. Plasmids and cloning primers used in this study are listed in Supplementary Table S2 and S3.

### Toxicity assays

BL21 cells were used for expression studies where only pBAD-derivative plasmids were tested while BL21(DE3) cells were used for co-expression studies from pBAD- and pET-derivative plasmids. Cells were transformed with plasmids and liquid cultures were inoculated and then grown to mid-late exponential phase (OD_600_ of ∼0.5-1). Cell suspensions of OD_600_=0.5 were prepared and 4 µL of serial 1:10 dilutions were spotted onto LB agar plates containing the appropriate antibiotics and either 0.8% (^w^/_v_) glucose or 0.4% (^w^/_v_) L(+)-arabinose and if appropriate IPTG for induction. For toxicity assays using anhydrotetracycline (aTc), cells were scraped directly from plates using a 1 µL loop, resuspended in LB and then directly streaked onto LB plates containing 0, 1, or 10 ng/mL aTc. Experiments shown are representative for at least two biologically independent repeats.

### Phage plaque assays

*E. coli* strains transformed with appropriate plasmids were grown to late exponential-early stationary phase and then diluted 1:20 into 0.35% LB soft agar kept liquid at 50°C. Appropriate antibiotics were added before pouring the soft agar onto LB plates containing the likewise antibiotics (ratio of 1:5 LB soft agar to LB agar). Phages were serially 1:10 diluted in phage buffer (10 mM Tris-HCl pH 7.4, 10 mM MgSO_4_, 0.1% (^w^/_v_) porcine gelatine) and 3 µL of the phage suspensions were spotted onto the solidified soft agar containing the *E. coli* strains. Plates were incubated at 30°C overnight (o/n) and plaques imaged the next day. Abortive infection phenotypes were likewise prepared, but only a concentrated sample of phage suspension was added and imaged repeatedly over 100 h. For isolation of Mav-infected *E. coli*, 50 mL LB medium was supplemented with 2% (^w^/_v_) gelatine and *E. coli* were diluted in therein to a final OD_600_=0.5 while Maverick phage suspension was added at an MOI of 10. Gelatine was allowed to set at 4°C, after which the tubes were incubated in a 30°C water bath for 45 min. The temperature was then increased to 37°C to completely melt the gelatine, followed by centrifugation to pellet the cells (5,000 × *g*, 5 min, 20°C).

### Protein purification

Rosetta strain BL21(DE3) was transformed with pNIC28-Bsa4 constructs encoding *E. coli* DarH, DarT3, ARH, ARH^1–442^, UvrD or *B. subtilis* DarT4, ARH and grown at 37°C in Terrific Broth supplemented with 50 μg/mL of kanamycin and 35 μg/mL of chloramphenicol. After reaching an OD_600nm_ of 1.0-1.2, the temperature was lowered to 18°C prior to induction of protein expression o/n by adding 0.5 mM IPTG. Harvested cells were resuspended in lysis buffer (50 mM HEPES (pH 7.4), 500 mM NaCl, 5 mM MgCl_2_, 5% glycerol, 20 mM imidazole, 0.5 mM TCEP. Then, cell suspensions were supplemented with Tergitol 15-S-9 of a final concentration of 0.8%, incubated at room temperature for 30 min on a rotating and then stored at -80°C until purification.

For protein purification, pellets were gently thawed and benzonase nuclease was added from in-house stock to a final concentration of 2.0 µg/mL. After incubation of 15 min at 4°C, cells were lysed by sonication and suspensions centrifuged for 45 min, 35000 x *g*, 10°C. Proteins were purified by immobilised metal affinity chromatography (IMAC) using Ni-Sepharose resin (Macherey-Nagel) and eluted stepwise in binding buffer containing 40–500 mM imidazole. Typically, a high salt wash with 1 M NaCl was combined with the first elution step including 40 mM imidazole. *E. coli* DarH, DarT3 and ARH^1-442^ were further purified by rebinding to Ni-IMAC after removal of the hexahistidine tag by recombinant TEV protease during o/n dialysis into buffer without imidazole. All proteins were finally purified by size-exclusion chromatography (SEC) (Superdex 75, GE Healthcare) in a buffer consisting of 25 mM HEPES (pH 7.5), 200 mM (*E. coli* ARH^1-442^)/250 mM (*E. coli* DarT3)/ 300 mM NaCl, 5% glycerol and 0.5 mM TCEP and supplemented with 5 mM MgCl_2_ for helicase and ARH enzymes.

All proteins were characterised by SDS-PAGE, then flash frozen in liquid nitrogen and stored at −80°C until required. Protein concentrations were determined by measuring absorption of the sample at 280 nm with the NanoDrop ND-1000 (Thermo Scientific).

### Analytical size exclusion chromatography

DarG_A_ complex studies by size exclusion chromatography were performed in buffer consisting of 25 mM HEPES (pH 7.5), 200 mM NaCl, 2 mM MgCl_2_, 5% glycerol and 0.5 mM TCEP. *E. coli* ARH full-length and ARH^1-442^ were diluted in assay buffer to a final concentration of 50 µM. 100 µL were injected onto a S200 10/300 equilibrated in assay buffer and fractions collected for SDS-PAGE analysis.

### Gel-shift assays for *in vitro* ADP-ribosylation activity testing

Oligonucleotide DNA substrates were purchased from Merck and Life Technologies-Invitrogen, RNA substrates from IDT. Sequences are listed in Supplementary Table S4. Oligo ADP-ribosylation reactions were performed in buffer containing 50 mM Tris-HCl pH 8.0, 50 mM NaCl (supplemented with 2 mM MgCl_2_ for assays performed with DarH and ARH) at 37°C for 60 min (30 min for motif preference of *E. coli* DarT3) in an assay volume of 10 µL. In general, 1 µM DarH/DarT protein was incubated with oligonucleotides at a concentration of 10 µM and 3 µM (dsDNA hairpin substrate) and 1 mM *β*-NAD^+^ and 2 mM ATP in presence of DarH. For following oligo de-modification by hydrolases, the ADP-ribosylation reaction was stopped by heating the samples for 15 min at 95°C. Then, samples were incubated with buffer as control or 1 μM of the indicated hydrolase at 37°C for 30 min. Reaction products were analysed by separation on denaturing polyacrylamide gels run in TBE buffer, after adding 10 µL urea loading dye (10 mM TRIS pH 8.0, 10 mM EDTA, 4 M urea) to the ADP-ribosylation reactions and incubation for 3 min at 95°C. 10 µL of the samples were loaded onto the gel and oligos visualised under UV light (340 nm) after SYBR Gold staining (Invitrogen) for 19 bp oligomers or SYBR Safe staining (Invitrogen) for dsDNA hairpin substrate.

### Immuno-Dot blot assays for detection of ADP-ribosylated DNA

BL21 cells transformed with pBAD33 plasmids (encoding DarH and DarT enzymes) and pNIC28-Bsa4 plasmids (encoding ARH enzymes) as indicated were grown to OD_600_ of 0.2-0.3 in LB containing 0.8% (^w^/_v_) glucose before protein expression was induced with 0.8% (^w^/_v_) arabinose and IPTG. Cells were harvested by centrifugation (4000 x *g*, 3 min), washed with PBS, re-suspended in boiling lysis buffer (1.0% SDS, 10 mM Tris, 1 mM EDTA, pH 8.0) and lysed by heating to 95°C for 5 min. Cell lysates were subjected to proteinase K treatment for 1 h, 50°C. gDNA was then extracted by phenol:chloroform:isoamyl alcohol (25:24:1) extraction and recovered by ammonium acetate/ethanol precipitation. The DNA pellets were washed twice with 70% ethanol before re-suspending in TE buffer and concentration determination using a NanoDrop ND-1000 (Thermo Scientific). 500-1000 ng of gDNA was dotted onto a nitrocellulose membrane (Amersham Protran 0.2 nitrocellulose) and crosslinked with 1200 J using a Stratalinker UV crosslinker. Crosslinked DNA was then immunoblotted for gDNA (autoanti-dsDNA, DSHB, 1:200) or ADPr-gDNA (Mono-ADP-Ribose, AbD43647, HRP- coupled, Bio-RAD) for 1h at RT in 5% (^w^/_v_) powdered milk in PBS-T. For detection of ADP-ribosylated DNA after Mav phage infection, membranes were incubated with an ADP ribose antibody o/n at 4°C. Secondary HPR-coupled antibody (anti-mouse, A16066, Invitrogen) was incubated 1:2000 in 5% (^w^/_v_) powdered milk in PBS-T at RT for 1 h. After antibody incubation steps, membranes were washed four times with PBS-T, before ECL-reagents were applied for ECL-based chemiluminescence detection using GE ImageQuant LAS 4000.

For DarH-DarT dependency and DarH cation and nucleotide preference testing, bacterial gDNA was extracted as described above. For DarH-DarT dependency, 10 µL assay reactions were set up with 500 ng/µL gDNA, 2 mM ATP, 2 mM MgCl_2_, 1 mM *β*-NAD^+^, 250 nM *E. coli* DarH, 250 nM *E. coli* DarT. For control reactions, ATP, or *β*-NAD^+^ were either omitted or MgCl_2_ replaced with 5 mM EDTA. Following an incubation for 20 min at 37°C, 2 µL of the samples were spotted onto 0.2 µm NC membrane as processed as described above. For cation preference testing, *E. coli* DarH was diluted to 2.5 µM in 10x assay buffer (500 mM Tris-Cl pH 8.0, 500 mM NaCl and 50 mM EDTA). Then, 10 µL assay reaction were set up with 300 ng/µL gDNA, 2 mM ATP, 250 nM *E. coli* DarH, 250 nM *E. coli* DarT and 25 mM cation in assay buffer. The reaction was initiated by addition of 1 µL of *β*-NAD^+^ to a final concentration of 1 mM after pre-incubation of the samples at 37°C for 2 min. 2 µL of the samples were spotted onto 0.2 µm NC membrane after 5 min, 10 min, 15 min. For nucleotide preference testing, *E. coli* DarH was diluted to 2.5 µM in 10x assay buffer (500 mM Tris-Cl pH 8.0, 500 mM NaCl). Then, 10 µL assay reactions were set up with 300 ng/µL gDNA, 2 mM MgCl_2_, 250 nM *E. coli* DarH, 250 nM *E. coli* DarT and 2 mM nucleotide in assay buffer. The reaction was initiated by addition of 1 µL of *β*-NAD^+^ to a final concentration of 1 mM after pre-incubation of the samples at 37°C for 2 min. 2 µL of the samples were spotted onto 0.2 µm NC membrane after 5 min, 10 min, 15 min.

### dsDNA unwinding assays

Cy5 and BHQ2-labelled oligos (IDT) were annealed in buffer consisting of 10 mM TRIS pH 8.0, 50 mM NaCl, 10 mM MgCl_2_ to a concentration of 10 µM. Unwinding assays were performed in assay buffer consisting of 20 mM HEPES pH 7.4, 40 mM NaCl, 5 mM MgCl_2_, 5% glycerol, 0.5 mM TCEP. First, 1:1 dilution series of UvrD and DarH with a top concentration of 2 µM were prepared in assay buffer and annealed DNA fork substrate diluted to 800 nM in assay buffer. Then, 10 µL of each protein dilution was added to 5 µL of DNA fork substrate (*i.e*. 200 nM final concentration) in black, low volume 384-well plates (Corning, #3820). The reaction was initiated by addition of 5 µL ATP of 4 mM (*i.e*. 1 mM final concentration), diluted in assay buffer from 100 mM stock solution. Fluorescence was immediately measured using a Spark microplate reader (Tecan) at 30°C, with excitation at 625 nm and emission at 665 nm, followed by measurements every 2.5 min for 60 min. For measurements in presence of DarT, DarH and DarT were prepared as dilution series together and the reaction was initiated by addition of 5 µL ATP/*β*-NAD^+^ of 4 mM (*i.e*. 1 mM final concentration). Samples were measured in triplicates and assays performed with two independent repeats.

### ATPase assays

ATPase activity was measured using a pyruvate kinase, lactate dehydrogenase enzyme linked absorbance assay in which the loss of absorbance at 340 nM (6,250 cm^−1^ M^−1^) is coupled to ATP hydrolysis. 80 µL reaction mixtures were set-up in triplicates in 96 well plates (Greiner UV-Star) and contained 1.0 U lactate dehydrogenase, 4.0 U pyruvate kinase, 0.2 mM NADH, 0.5 mM PEP, 1 µM DarH or 100 nM UvrD in 20 mM HEPES pH 7.5, 200 mM NaCl, 5 mM MgCl_2_. For measuring DNA- stimulated ATPase activity, the reaction mix contained in addition 2 μM of single stranded 27 bp DNA. To determine the assay window, high controls were prepared by omitting the helicase enzyme from the reaction, whereas low controls were prepared by replacing NADH with buffer. The reaction was initiated by the addition of ATP, and absorbance at 340 nm was measured using a Spark microplate reader (Tecan) at 37°C every minute for 20 min. Samples were measured in triplicates and the assay was performed with three independent repeats.

### Preparation of ADP-ribosylated guanines and adenine bases for LC-MS/MS analysis

20 µg of PolyT-A and PolyT-G oligonucleotides were ADP-ribosylated with *E. coli* DarT3, *B. subtilis* DarT4 and *S. coelicolor* ScARP, respectively, in assay buffer consisting of 50 mM Tris-Cl pH 8.0, 50 mM NaCl, 0.5 mM *β*-NAD at 37°C in a total volume of 60 µL. *E. coli* DarT3 was used in a 1:10 molar ratio to DNA, *B. subtilis* DarT4 in 1:2 molar ratio to DNA, and ScARP in a 1:100 molar ratio. Then, samples were treated with 100 U nuclease P1 (NEB), 10 U antarctic phosphatase (NEB) and 1 U phosphodiesterase I from *Crotalus adamanteus* venom (SVPD, Millipore Sigma, dissolved in 40 mM TRIS pH 8.4, 10 mM MgCl_2_, 200 mM NaCl) in buffer containing 25 mM TRIS pH 7.6, 50 mM NaCl, 1 mM ZnCl_2_, 10 mM MgCl_2_ for 4.0 h at 37°C.

### LC-MS/MS data acquisition

Data acquisition was performed using a Vanquish Flex UHPLC system (Thermo Fisher Scientific) coupled to a Q Exactive HF mass spectrometer (Thermo Fisher Scientific). Samples were diluted 1:10 in buffer A (0.1% formic acid), and 2 µL of each sample was injected onto a Luna Omega Polar C18 column (2.1 × 100 mm, 1.6 µm; Phenomenex) operated at a flow rate of 200 µL/min. Chromatographic separation was achieved using buffer B (100% acetonitrile, 0.1% formic acid), which was held at 0% for 2 minutes, increased to 95% over 12 minutes, and maintained at 95% for an additional 2 minutes. The column was subsequently re-equilibrated at 0% buffer B for 4 minutes. The mass spectrometer was operated in positive ionization mode with data-dependent acquisition. Full MS¹ scans were acquired over an m/z range of 85–700 at a resolution of 120,000. Heated electrospray ionization (HESI) source parameters were as follows: sheath gas (N₂) flow rate of 47, auxiliary gas flow rate of 10, and sweep gas flow rate of 2 (all in arbitrary units). The spray voltage was set to 3.20 kV, the capillary temperature to 250 °C, and the auxiliary gas heater temperature to 380 °C. The five most intense precursor ions were selected for fragmentation by higher-energy collisional dissociation (stepped HCD with normalized collision energies of 20, 40, and 60). MS² spectra were acquired at a resolution of 30,000. The automatic gain control (AGC) targets were set to 3 × 10⁶ for MS¹ and 1 × 10⁵ for MS², with maximum injection times of 200 ms and 50 ms, respectively. MS² spectra corresponding to the [M+H]⁺ precursor ions of ribosylated guanosine (theoretical m/z 400.1463; observed m/z 400.1456) and ribosylated adenosine (theoretical m/z 384.1514; observed m/z 384.1506) were extracted directly from the .raw files using a custom PowerShell script built on the Thermo RawFileReader library (Thermo Fisher Scientific). For each compound, the spectrum was taken as the MS² scan whose recorded precursor matched the target mass within 0.02 Th and which lay closest in retention time to the apex of the corresponding chromatographic peak, in every case within one duty cycle of it. Spectra were exported as centroided mass/intensity lists from the Orbitrap label data.

## Data analysis

Gene neighbourhood analysis was performed using webFlaGs (https://server.atkinson-lab.com/webflags) ^45^with *E. coli* DarT3 as the query sequence. Default settings were applied with the maximum number of BLAST hits set to 200. Predictions of the structures of individual DarHTG components and predications of ARH dimers and *T. ammonificans* ARH-Mav24 interactions were done using AlphaFold3^24^ with default settings. Sequence conservation mapping was performed using ConservFold^46^. Structural homology searches based on the AlphaFold3-predicted structures of DarHTG components were done using Foldseek^22^. Structural alignments and analyses, as well as figure preparation, were carried out using PyMol (Molecular Graphics System, Version 3.0 Schrӧdinger, LLC). For multiple-sequence alignments, JalView v2.11.5.2^47^ and MAFFT7^48^ was used. The phylogenetic tree for DarT enzymes was generated with SplitsTree4 (v4.19.2) using the Neighbour- Joining (NJ) method^49^ and confidence levels estimated using 1000 cycles of the bootstrap method. To sequence the Mav escaper mutants, approximately 11-12 kb pieces, each overlapping around 200- 300 bp with the next, where PCR amplified using primer pairs RedMav-1/RedMav-2, RedMav- 3/RedMav-4, RedMav-5/RedMav-6, and RedMav-7/RedMav-8, respectively. Fragments were sent for Oxford Nanopore sequencing and reads assembled using SnapGene v8.2.2. GraphPad Prism v10 was used for data visualisation, statistical analysis and plotting. Inkscape v1.4.2. was used for figure preparation. ChemDraw v26 was used for analysis and drawing of LC-MS/MS fragments generated from ADP-ribosylated guanine and adenine.

## Acknowledgments

We thank Professor Elena Conti and her team for the helpful discussions. Work in the Marion Schuller Laboratory is supported by the Lise-Meitner Excellence Programme of the Max Planck Society. We thank the Mass Spectrometry facility of the MPIB (RRID:SCR_025745) for the performance of the LC-MS/MS analysis. The work in Ivan Ahel’s lab has been supported by the following grants: Biotechnology and Biological Sciences Research Council (BB/R007195/1 and BB/W016613/1), Wellcome Trust (210634, 223107, and 302632), CRUK (C35050/A22284), and the Guy Newton Fund. We thank Professor Tim Blower for providing a copy of the Durham Phage Collection^21^ and Professor Alexander Harms for providing a copy of the BASEL collection^31,32^ and host strains.

## Author contributions

MS conceived the project and conceptualised experiments with input from RL and IA. RL conducted cloning, protein expression/purification, *in vivo* assays. MS conducted bioinformatic, biochemical and structural studies. ALG, JF, YW, JH, JZ, SG, YL, AM assisted with protein purifications, biochemical and *in vivo* assays; OS cloned pBAD33 control vector; YZ, CT supported performance of phage assays. RL and MS wrote the manuscript with support of all other authors.

## Declaration of interests

The authors declare no competing interests.

## Data availability

All data supporting the findings of this study are available within the manuscript and its Supplementary Information. All other data generated and analysed as well as study-specific materials generated during this study are available upon request to the corresponding author.

## Notes

### Competing Interest Statement

The authors have declared no competing interest.

